# Query Augmented Generation (QAG) from the Genomic Data Commons for Accurate Variant Statistics

**DOI:** 10.1101/2025.09.02.673797

**Authors:** Aarti Venkat, William P. Wysocki, Michael Lukowski, Steven Song, Anirudh Subramanyam, Zhenyu Zhang, Robert L. Grossman

## Abstract

In precision oncology, researchers often use public knowledgebases to check somatic variant frequencies against their cohort data. Large language models (LLMs) can quickly answer questions on somatic variant frequencies, but often hallucinate and give inaccurate results for factual data. Using synthetic queries, we show that somatic variant frequencies in baseline LLM responses are underestimated compared to the Genomic Data Commons (GDC), the world’s largest data commons for cancer research. We present a modular architecture called Query Augmented Generation (QAG) for integrating LLMs with high-quality data from a third party data source such as a data commons, knowledgebase or database. We apply QAG to the GDC to help researchers obtain accurate frequencies for somatic variants, copy number variants, and MSI status—even for complex queries requiring multiple steps in the GDC portal and API. Our software is deployed as a model context protocol (MCP) server on Hugging Face and available on GitHub.

## 1 Introduction

In precision oncology, frequencies of somatic or germline mutations, copy number variants, and microsatellite instability status are typically estimated for every patient from sequencing data and compared with frequencies from public knowledgebases, databases or data commons to understand mutation prevalence and inform clinical decisions [1–3]. Knowledgebases such as the Genomic Data Commons (GDC) provide an excellent resource for accessing single variant frequencies through the portal; however, more complex queries involving combinations of variants often require a multi-step process and familiarity with the GDC API for programmatic execution [4, 5]. To help researchers obtain quick answers to such queries, we present an architecture called Query Augmented Generation (QAG) that integrates Large Language Models (LLMs) with third party data sources such as knowlegebases, databases and data commons (Figure 1). In QAG, a natural language query from a user is decomposed into relevant entities and a “query intent” is inferred to build a set of query or queries that can be executed against a third party data source. The result of this execution is used to build LLM prompts and confirm the accuracy of the LLM response (Figure 1). In this study, we use the GDC as the external data source and prototype QAG for queries in cancer research. Using functionality wrapped around the rich collection of API endpoints in the GDC, the GDC-QAG algorithm provides accurate answers to simple and complex queries concerning frequencies of simple somatic mutations (SSMs), copy number variants (CNVs), combination variants and frequency of microsatellite instability (MSI).

**Figure 1:**
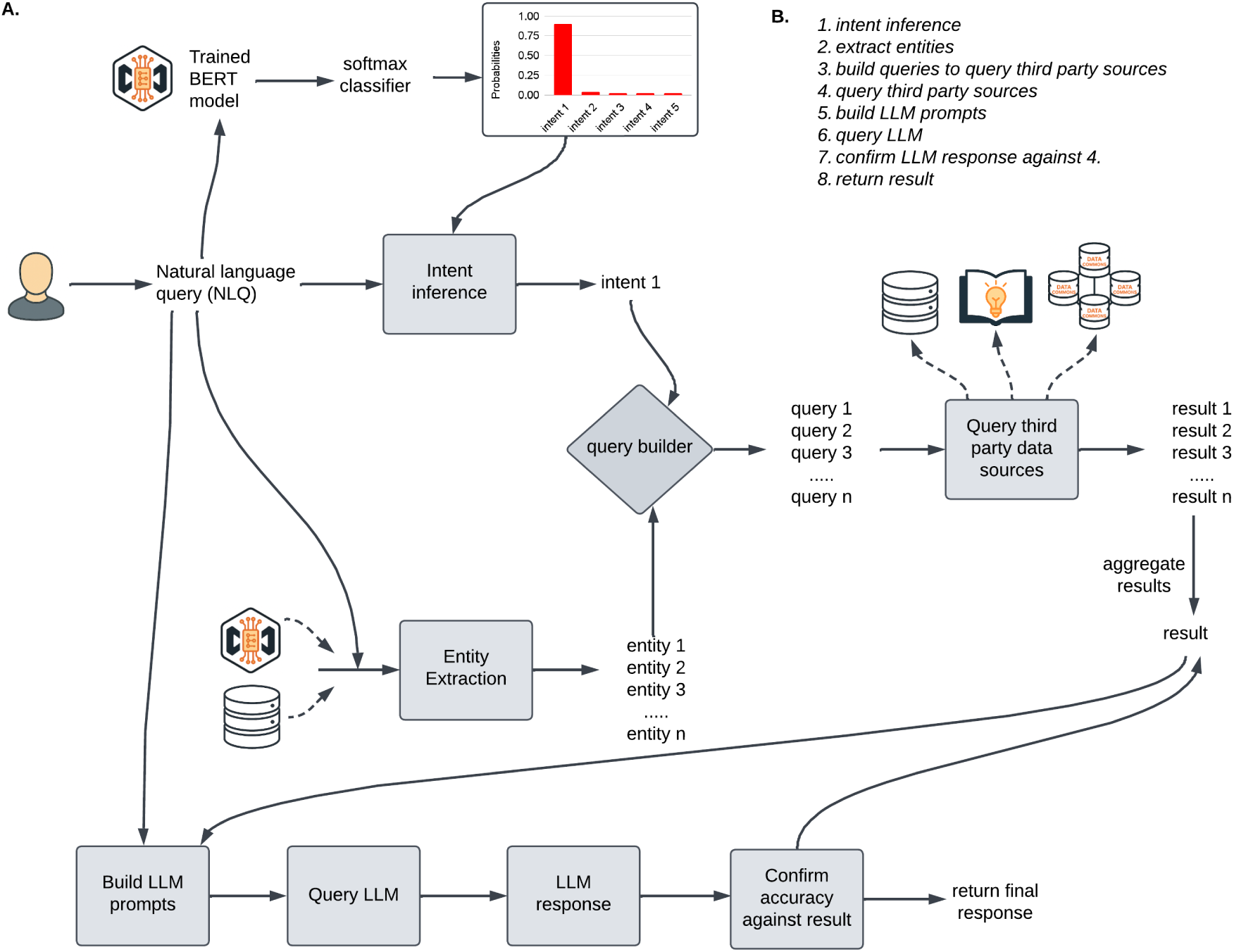
Query Augmented Generation (QAG) architecture for integrating LLMs with third party data sources such as databases, knowledgebases and datacommons. A. Grey boxes represent the steps in QAG. A natural language query from a user is passed through an *Intent inference* step to extract a “query intent” using a trained model, which helps map the user query to a relevant API call. The *Entity Extraction* step extracts relevant entities from the query using information from an external source such as a database. A trained model could also alternately be used. For biomedical queries, entities can be genes, mutations, disease such as cancer types. Depending on the intent, a query or queries are built to query a third party source such as a database, data commons or a data mesh. Queries are a collection of API calls. A result for each query is obtained and aggregated to obtain one result. This result, together with the input user query, is used to build LLM prompts and query the LLM. The LLM response is checked against the result obtained from querying third party data sources to ensure accuracy, and returned as the final response. B. The steps in A. are outlined. All logos are generated using the Microsoft M365 Copilot app.

The GDC is a gold-standard knowledgebase that offers quick insights into single somatic variant frequencies through the data portal, for researchers to compare with variant frequencies estimated from their patient cohorts. Complex queries can also be executed manually through the portal, but this often involves multiple steps and is time consuming. For example, if a researcher is interested in the frequency of cases carrying deep deletions in both CDKN2A and CDKN2B in mesothelioma, they can do so through the GDC portal using multiple apps in multiple steps: First, they need to create a cohort for TCGA-MESO using cohort builder, then create a mutated gene filter for CDKN2A and CDKN2B, then save two cohorts for CDKN2A and CDKN2B homozygous deletions and select them using the set operators app to uncover a joint frequency of 44.83%. In another example, if a researcher is interested in the frequency of cases carrying both IDH1 R132H and TP53 R273C mutations in lowgrade glioma, they would similarly need to create a cohort of patients with low-grade glioma, upload the two somatic mutations in the mutation frequency app, create two cohorts: one for IDH1 R132H and the other for TP53 R273C, and finally use the two cohorts in the set operations app to uncover a joint frequency of 7%. If a researcher has multiple such queries, it can be manually laborious to do so through the portal. As an improvement over the GDC portal, the GDC-QAG algorithm can be particularly useful for obtaining frequencies of cases carrying mutation combinations.

QAG is an architecture that integrates an LLM with a knowledgebase like the GDC to generate a response after query augmentation (Figure 1). Recent advances in LLMs are transforming the way scientific research is conducted [6–13]. These models offer ideas and solutions through simple conversations involving natural language queries, making them potentially suitable to gain quick answers to cancer research questions surrounding specific mutations and frequencies. Previous publications have noted that hallucinations are often more pronounced for factual information involving numeric or statistical facts, which can now be mitigated by frameworks such as Retrieval Augmented Generation (RAG), Retrieval Interleaved Generation (RIG) and Model Context Protocol (MCP) [14–18]. Through these frameworks, a knowledgebase or tooling around a knowledgebase can be connected to an LLM to obtain accurate responses to natural language user queries, including programmatic execution of more complex queries involving multiple API calls. The GDC-QAG algorithm is containerized using Gradio and deployed as an MCP server app on Hugging Face. Accurate context provided to the LLM, together with a confirmation step, helps to ensure the final output from QAG is accurate (Figure 1).

### 1.1 Contributions of this study

The main contributions of our study are listed below:

#### 1.1.1 The QAG architecture enables integration of an LLM with a knowledgebase for biomedical research

QAG is an approach to integrate high-quality data from a biomedical knowledgebase, database or data commons with an LLM at inference time to generate trustworthy responses to user queries. Our study applies QAG to questions of biomedical importance, specifically cancer research. Applied to the GDC, researchers who lack knowledge of GDC APIs can easily run the GDC-QAG algorithm, and can do so in resource-limited settings without elaborate infrastructure. QAG is prototyped on the llama-3B LLM, but is modular to any choice of LLM.

#### 1.1.2 Assess the degree of discrepancy between variant frequencies in baseline LLM responses and those found in the GDC

Several studies have explored the use of LLMs in genomics, but to our knowledge, there is no quantification of the accuracy of somatic variant frequencies in baseline LLM responses for different precision oncology queries [13, 19–25]. Using curated data on somatic variants from the CIViC database, the GDC, and previous publication, we constructed an evaluation dataset called the *GDC-Variant-Testset* (*GV T*), consisting of n=6011 synthetic queries spanning several use cases in precision oncology, together with their paired variant frequencies obtained from the GDC (Methods) [4, 5, 10]. We examined the difference between the variant frequencies in three baseline LLM responses and *GV T* : gpt-4o-2024-08-06, meta-llama/Llama-3.2-3B-Instruct and Qwen/Qwen1.5-4B-Chat, herein referred to as GPT-4o, llama-3B and Qwen-4B, respectively. These models were chosen to represent a mix of widely-recognized closed-source (GPT-4o) and open-source models (Qwen-4B, llama-3B) and model sizes. The smaller models were chosen given their previous performance in various benchmark tasks related to reasoning, tool use, math and long context, to compare with a large model like GPT-4o for precision oncology queries, and their ease of use for academic research and testing [26, 27]. We also evaluated GDC-QAG on *GV T* and demonstrate that it produces accurate responses, whereas variant frequencies in baseline LLM responses are underestimated compared to the GDC.

#### 1.1.3 Trained BERT classifier infers the appropriate GDC API endpoint to execute from a natural language query

The GDC-QAG algorithm uses a BERT classifier trained on a diversity of paired natural language queries and query type (“intent”) to help map a query to appropriate GDC API endpoint(s). For example, the classifier will predict an *ssm frequency* intent for queries concerning frequencies of SSMs, which then lead to building and execution of *ssm frequency* specific API calls.

#### 1.1.4 The GDC-QAG algorithm and its implementation enable programmatic execution of complex queries that are not available in the GDC today

Using GDC-QAG, researchers can simply provide an input file with questions spanning combinations of SSMs and CNVs and obtain variant frequency results. GDC-QAG leverages several GDC API endpoints to calculate joint frequencies of variants, thereby providing a single tool to enable this computation. This information can be used to systematically investigate epistatic and other functional interactions between somatic mutations in the protein, build a mutation frequency profile of combinations of mutations, and even discover new variant combinations that improve or worsen survival outcomes. Such information, that is not widely available in publications but niche to the GDC is especially valuable to improve LLM responses using GDC-QAG.

## 2 Results

### 2.1 Shortcomings in ChatGPT for cancer genomic queries

Our investigation of select queries spanning frequency of a single SSM, CNV and joint frequencies of SSMs or CNVs showed that the data retrieved by ChatGPT appears often to be associated with a wrong gene, variant type, cancer type or cohort, than that stated in the query (Table S1, Methods). Our findings are consistent with previous publications that urge researchers to manually verify the references with the generated response [28–30].

### 2.2 Variant frequencies in baseline LLM responses are underestimated compared to the GDC

To more comprehensively quantify the difference between variant frequencies in baseline LLM responses and frequency in the GDC, we constructed *GV T* using data from different sources: i) data from curated clinical evidence nightly summaries available in the CIViC database, ii) data from the GDC, and iii) data from a previous publication focused on pancancer homozygous and heterozygous deletions [31] (Methods). Briefly, we generated synthetic template questions with placeholders for gene, mutation and disease that map to different precision oncology use cases and used molecular information from the different sources to curate *GV T* (Tables S2-S5). Corresponding variant frequencies were obtained by independently querying the GDC API using relevant parameters (Methods).

Overall, the *GV T* comprises expert curated data from 115 unique genes and 54 somatic mutations across 62 different cancer types. We divided the queries in *GV T* into five different query types or intents: i) queries pertaining to frequencies of cases with SSMs (*ssm frequency*) (n=144), ii) queries pertaining to the frequency of cases with SSMs or CNVs, or both (*cnv and ssm*) (n=4206), iii) queries pertaining to the frequency of CNV losses or gains (*freq cnv loss or gain*)(n=1453), iv) queries pertaining to frequency of MSI (*msi h frequency*) (n=38), and v) queries pertaining to the frequency of combinations of CNV losses or gains (*freq cnv loss or gain comb*) (n=170). Example questions for each intent are shown in Table 1.

**Table 1:** Example questions by intent. The sample size for each intent (*GV T* Count) reflects the number of queries in *GV T* that satisfy the ground-truth frequency cut-off of atleast 5%, and the sampling process for curation of *GV T* (Methods). For instance, for the *ssm frequency* intent, there are only a handful of simple somatic variants (*≈* 0.002%) in the GDC whose frequency is *≥*5% in at least one project. MSI frequency queries test the frequency of MSI in different cancers.

| Intent | Example Question | *GVT* Count |
| --- | --- | --- |
| MSI frequency | In Colon Adenocarcinoma TCGA-COAD project, what is the occurrence rate of microsatellite instability in the genomic data commons? | 38 |
| Frequency of CNV loss or gain | What is the frequency of somatic JAK2 heterozygous deletion in Acute Lymphoblastic Leukemia - Phase II TARGET-ALL-P2 project in the genomic data commons? | 1453 |
| CNV and SSM | What is the incidence of simple somatic mutations or copy number variants in PDGFRA in the genomic data commons for Uterine Carcinosarcoma TCGA-UCS project ? | 4206 |
| SSM frequency | What is the rate of occurrence of NRAS Q61R mutation in Skin Cutaneous Melanoma TCGA-SKCM project in the genomic data commons? | 144 |
| Frequency of CNV loss or gain (combinations) | What is the co-occurrence frequency of somatic heterozygous deletions in CDKN2A and CDKN2B in the mesothelioma project TCGA-MESO in the genomic data commons? | 170 |
|  | *GVT* Total | 6011 |

We defined a measure *δ*, as the difference between variant frequency in baseline LLM response and *GV T* . We performed baseline evaluations of three LLMs and GDC-QAG using reproducible parameters, prompting the models to only use data from the GDC (Methods). Importantly, these evaluations are done with zero shot prompting, so the models rely solely on information seen during their training. Figure 2 shows that on average, the variant frequencies in baseline LLM responses are underestimated compared to that in the GDC often with high variability and outliers. The histograms of variant frequency differences display a broad distribution (Figures S1,S2, S3). For the *cnv and ssm* intent, a median negative *δ* of *≈ −*26% is obtained for both llama-3B and GPT-4o (Figure 2). Qwen4B also performed similarly, with a median *δ* of about -22%. For example, the query “What is the incidence of simple somatic mutations or copy number variants in PDGFRA in the genomic data commons for Uterine Carcinosarcoma TCGA-UCS project?” returned a llama-3B model response of 0.03%, GPT-4o response of 5% and Qwen-4B model response of 10.4%. The frequency in the GDC is 47.37%, resulting in a very negative *δ* of -47%, -42% and -36%, respectively. For the limited queries tested, the GPT-4o model appears to return frequencies that are closer to the GDC for the *ssm frequency* intent than other models (Figure 2).

**Figure 2:**
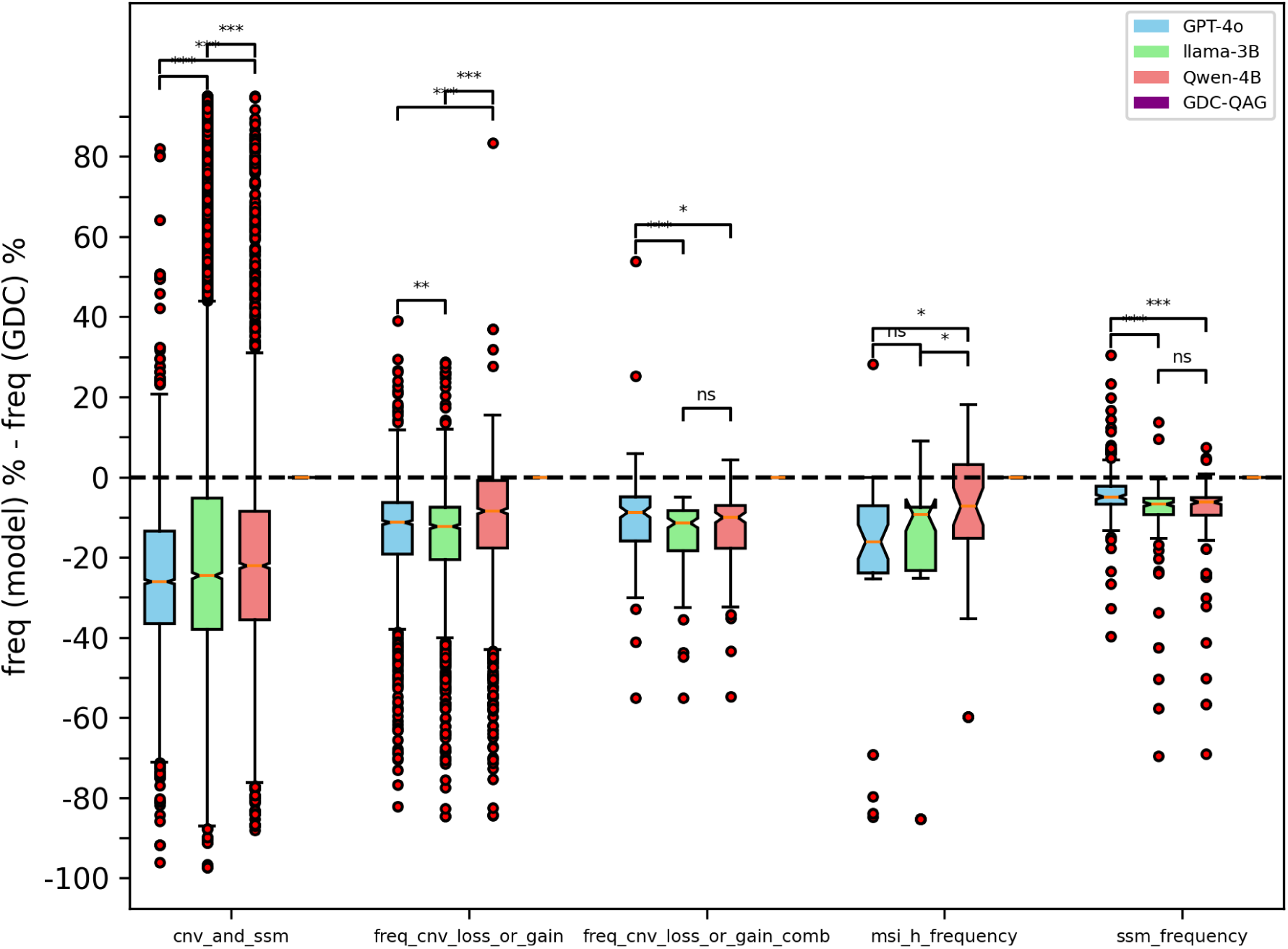
This plot shows the difference in variant frequencies between model generated responses and that of the GDC API. Results are grouped by five different use cases. Black dotted line points to no difference between model frequencies and that from the GDC. GDC-QAG variant frequencies match up exactly with the GDC API. Significance calculated using t-tests. **\*\*\*** *P <* 0.001, **\*\*** *P <* 0.01 , **\*** *P <* 0.05, **ns** *non − significant*

The intent *freq cnv loss or gain* is composed of queries containing heterozygous deletions, gain or amplification frequencies in different genes. To learn if specific categories of questions cause the models to perform worse than other categories, we stratified LLM performance by the type of CNV: heterozygous deletions (n=819), gains (n=642) or amplifications (n=160) (Figure 3). Both llama-3B and GPT-4o models underestimate frequencies for all types of CNVs. The Qwen-4B model has a very fast runtime and overall returns frequencies that are closer to the GDC for ‘amplification’ and ‘gain’ queries compared to the other two LLMs (Figure 3). We also tested if queries from TCGA projects lead to better overall performance by the LLMs than queries from non-TCGA projects, given the preponderance of publications from TCGA projects. We found no performance improvement specific to TCGA project queries across LLMs and intents (Figures S4,S5, S6).

**Figure 3:**
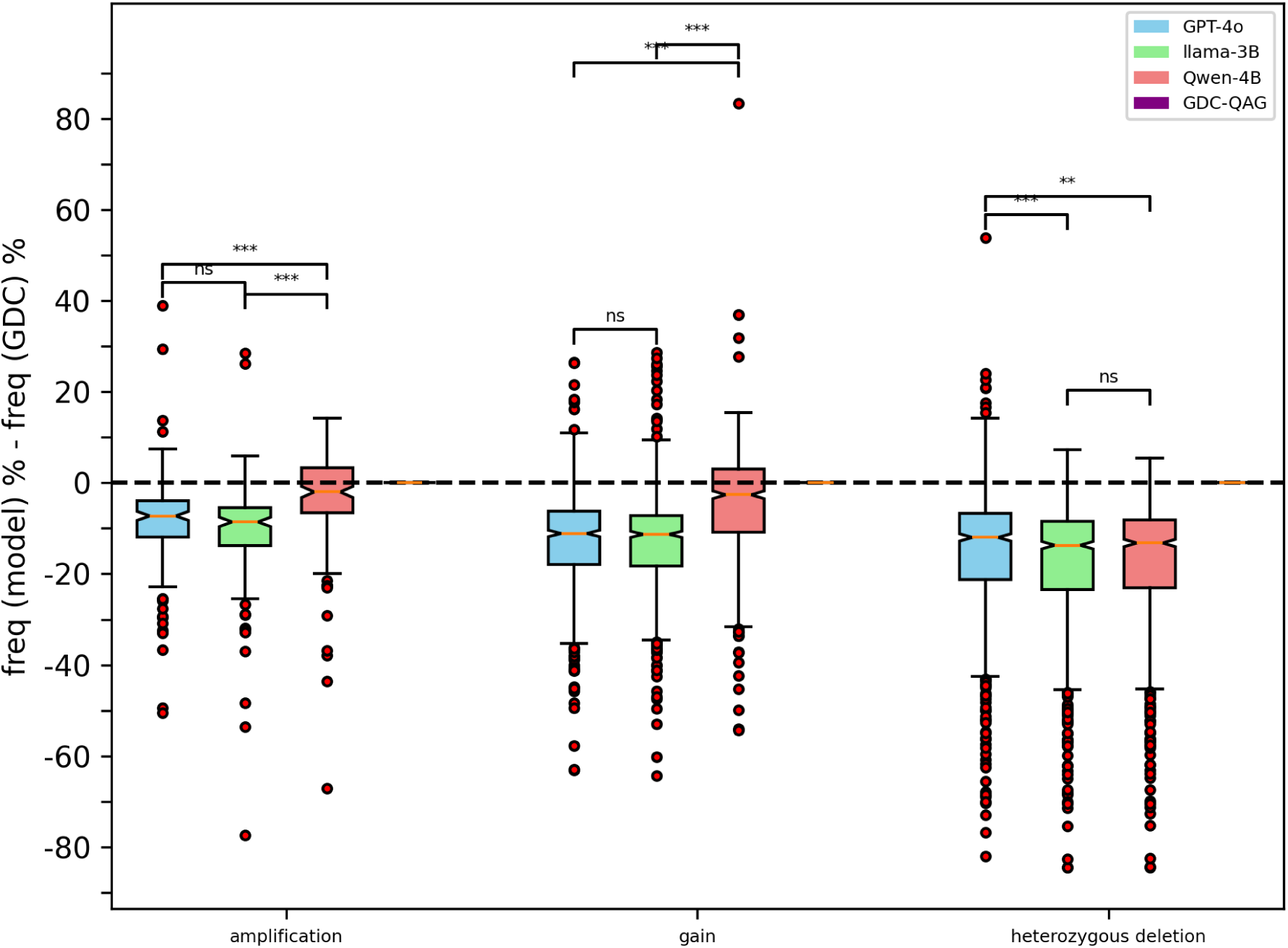
This plot shows the difference in variant frequencies between model generated responses and that of the GDC API for different types of copy number variants. Black dotted line points to no difference between model frequencies and that from the GDC. GDC-QAG variant frequencies match up exactly with the GDC API. Significance calculated using t-tests. **\*\*\*** *P <* 0.001, **\*\*** *P <* 0.01 , **\*** *P <* 0.05, **ns** *non − significant*

### 2.3 GDC-QAG generates accurate variant frequencies on ***GV T***

We evaluated the accuracy of GDC-QAG on *GV T* and found complete concordance between the variant frequencies produced by GDC-QAG and *GV T* (Table 2). Figures 2 and 3 further illustrate that the variant frequencies from GDC-QAG are identical to those from the GDC API, with no discrepancies.

**Table 2:** % agreement is calculated by comparing GDC-QAG variant frequencies with that of *GV T* . *GV T* frequencies are generated by dividing the synthetic queries into seven different query types as shown in the “Query type” column. For each query type, the GDC API is queried using independent functions that accept the relevant GDC API parameters and endpoints as indicated, and *GV T* frequencies generated.

| Query type | *GVT*<br>Count | API parameters | endpoints | % agreement<br>(GDC-QAG<br>and *GVT*) |
| --- | --- | --- | --- | --- |
| CNV and SSM | 4206 | gene<br>project | /cnv_occurrences<br>/ssm_occurrences<br>/case_ssms | 100 |
| Heterozygous<br>or Homozygous<br>deletions | 651 | gene<br>project<br>Loss or Homozygous-Deletion | /cnv_occurrences<br>/case_ssms | 100 |
| Gains | 642 | gene<br>project<br>Gain | /cnv_occurrences<br>/case_ssms | 100 |
| Amplifications | 160 | gene<br>project<br>Amplification | /cnv_occurrences<br>/case_ssms | 100 |
| SSM frequency | 144 | gene<br>project<br>protein mutation | /ssm_occurrences<br>/case_ssms | 100 |
| MSI frequency | 38 | project | /files | 100 |
| Combination<br>heterozygous<br>or homozygous<br>deletions | 170 | genes<br>project<br>Loss or Homozygous-Deletion | /cnv_occurrences<br>/case_ssms | 100 |

Overall, the results presented in Figures 2 and 3 show that to increase confidence and trustworthiness in variant frequencies reported by LLMs, the query could be augmented with variant frequencies from an external data source and used by the LLM for a more accurate response generation. All baseline evaluation results are available in Supplementary Tables S6-S9.

### 2.4 GDC-QAG algorithm: steps and implementation

The GDC-QAG algorithm is an application of the QAG architecture to the GDC (Figure 1). Two use cases surrounding frequencies of cases with multiple somatic mutations and copy number variants are presented (Figures 4, 5). The individual steps are described below.

**Figure 4:**
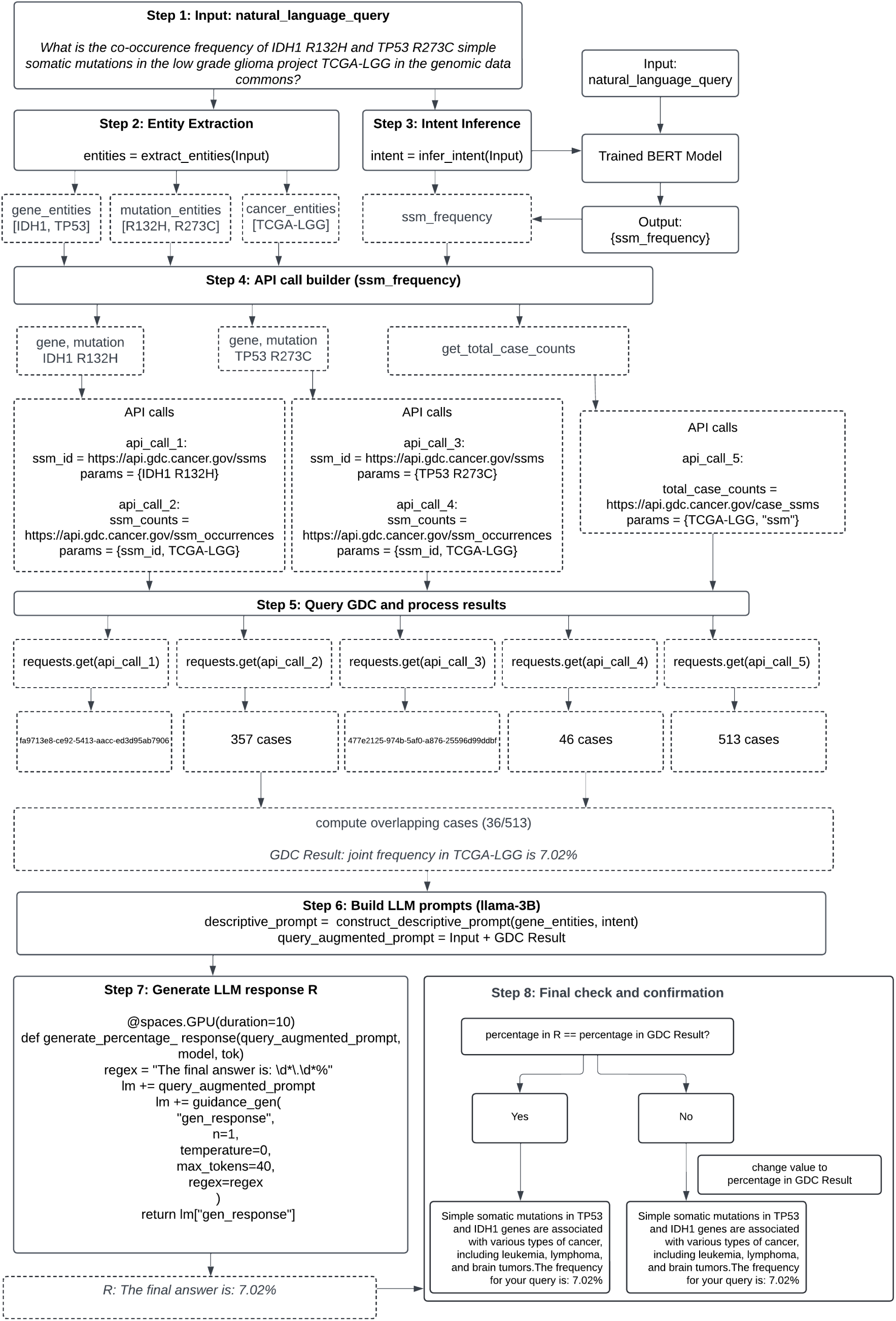
GDC-QAG algorithm, focusing on a combination somatic mutation use case from the GDC. Solid boxes represent the steps, and dotted boxes represent the output of the steps. Step 1: An example query concerning the frequency of a combination of SSMs in low grade glioma is passed as the input. In Step 2, the entity extraction step extracts gene, mutation and cancer entities from the query. In Step 3, a trained BERT classifer model predicts an intent label from the query, in this case *ssm frequency*. In Steps 4 and 5, for the gene mutation pair in question, five different API calls will be built, executed and processed into a text-formatted GDC Result with the joint frequency. In Step 6, LLM prompts are built. A descriptive prompt is constructed to obtain a natural language response from the LLM based on genes and query intent. A query augmented prompt is also built, using the GDC Result appended to the query from Step 1. In Step 7, the query augmented prompt is passed to the llama-3B model and an LLM response R (variant frequency) is extracted using constrained decoding. In the same step, the descriptive prompt is also passed to the llama-3B model using a similar set up, to extract a descriptive natural language response using constrained decoding. A final confirmation step checks the LLM response R to ensure accuracy of the variant frequency, and a QAG response is returned by concatenating the descriptive natural language response and the variant frequency. The Gradio app with the GDC-QAG algorithm is configured as an MCP server.

**Figure 5:**
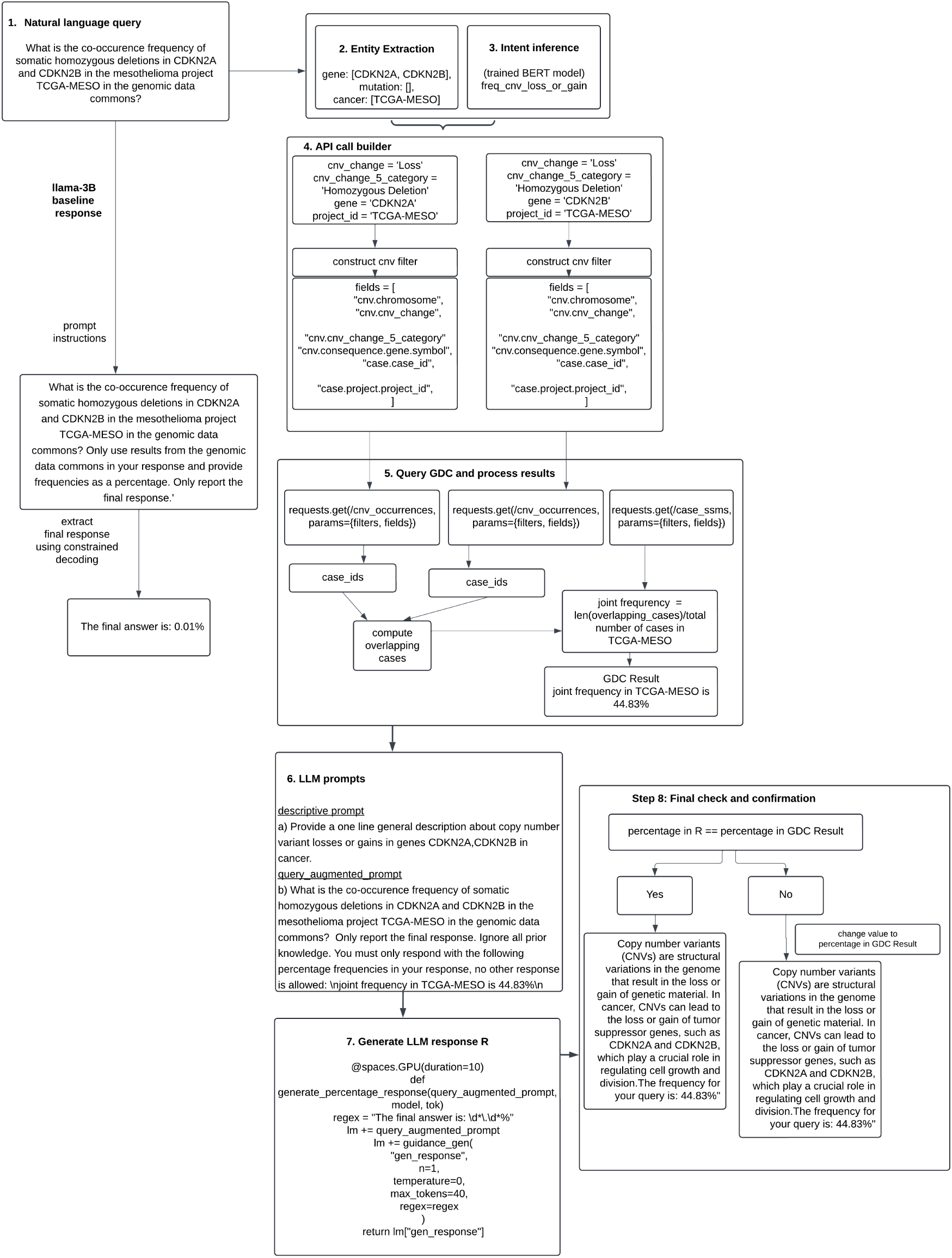
GDC-QAG algorithm, focusing on a combination copy number frequency use case. For the natural language query passed as input, CDKN2A and CDKN2B genes are infered as gene entities, cancer entity is the TCGA-MESO project for mesothelioma. The *freq cnv loss or gain* intent is output by the trained BERT classifier model in Step 3. Using the entities and intent, API calls are built using the /cnv occurrences endpoint in Step 4. In Step 5, one API call is executed for each gene and a case ids list is obtained, with the list of cases carrying the copy number variant. A list of shared cases is then computed, and counts are converted into frequency using a third API call to /case ssms endpoint and a formatted GDC Result is generated with the joint frequency. A query augmented prompt is generated with the GDC Result in Step 6, LLM prompts are built and responses generated using constrained decoding in Step 7 using the same format as shown in Figure 4. The final confirmation step ensures the variant frequency in the final response matches the frequency obtained in step 5 , which is 44.83% in this case. The Gradio app with the GDC-QAG algorithm is configured as an MCP server.

#### 2.4.1 Entity Extraction

The entity extraction step consists of decomposing a query into the relevant entities for constructing the required GDC API call parameters. The entities in question are genes, protein mutations and cancer types. The genes and protein mutations are inferred from the query using a simple dictionary-based look-up of terms in the query that map to keys and values in a dictionary constructed from GDC data (Methods). For identification of specific cancer types from the query, we integrated a combination of different methods that use the /projects GDC API endpoint and scispacy NLP models from allenai [32] (Methods). The goal of cancer entity extraction is to infer cancer entities in the form of a GDC project id such as TCGA-BRCA, that enables an easy construction of the GDC API call.

#### 2.4.2 Intent Inference

In the Intent inference step, an intent label is predicted from the query. The intent label infers a query intent, which could be to obtain SSM frequency, CNV frequency, MSI, or CNV and SSM frequency. The intent label maps the query to appropriate GDC API call(s) to be executed. To achieve this, we trained a BERT classifier using paired synthetically generated training data containing queries and labeled intents (Figure 6, Methods). The trained BERT classifier outputs an intent label for the natural language query such as that shown in Table 1, which guides subsequent steps for API call building and execution.

**Figure 6:**
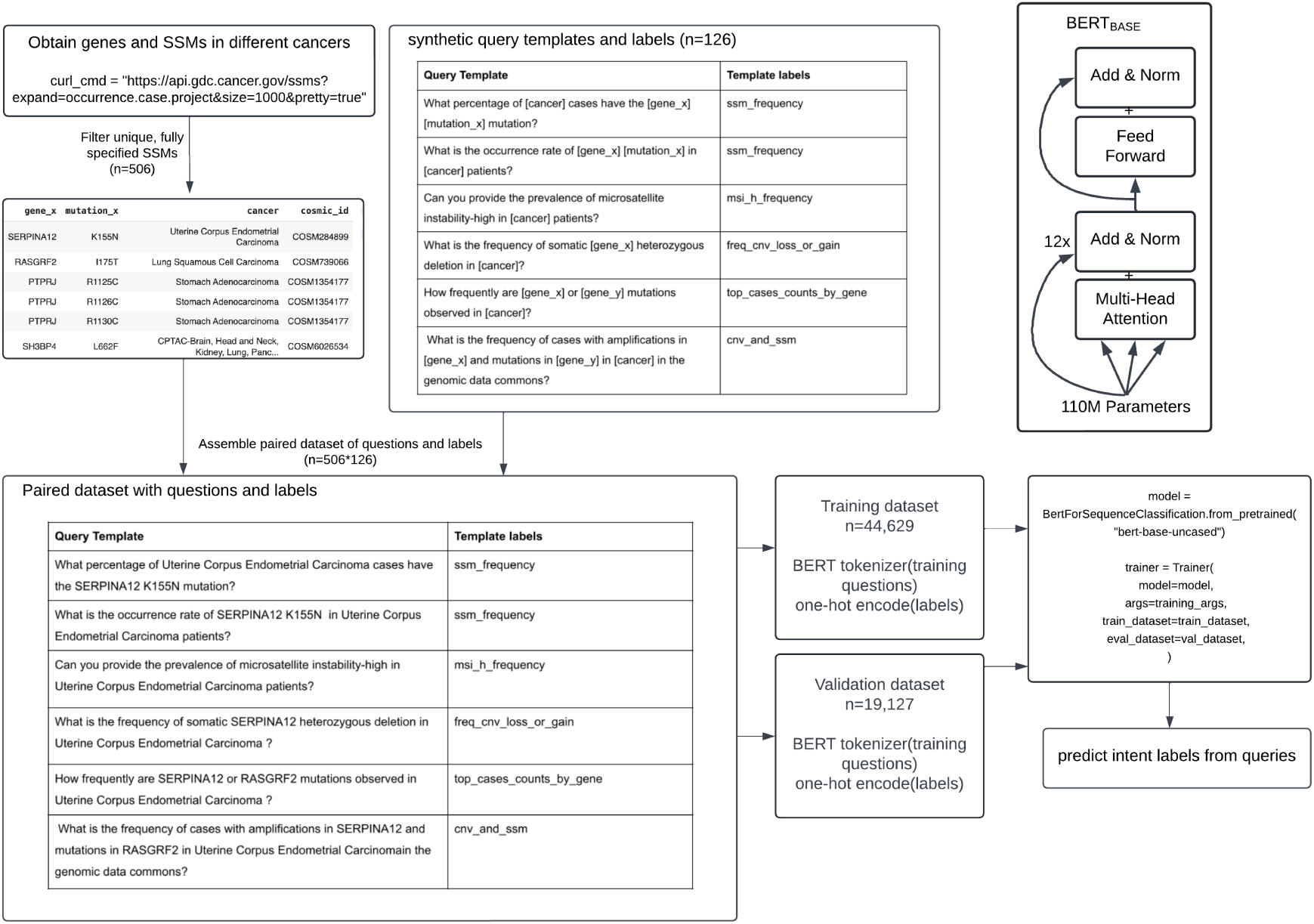
A *bert-base-uncased* classifer model is trained to infer intent from a query. A synthetic dataset is assembled with paired questions and labels using data from the /ssm endpoint in the GDC (n=63756). A snapshot of output from /ssm endpoint is shown, along with query template label pairs and the synthetic dataset. The dataset with 63756 rows is split into training and validation datasets (70:30 split) and questions tokenized using the *BertTokenizer*, labels one-hot encoded. Training and validation datasets consisting of tokenized questions and one-hot encoded labels were created using PyTorch and input into Trainer from the Transformers library for training. The trained model is used for intent prediction in GDC-QAG. Top-right, BERT size and architecture, the trained classifer model has 110M parameters. Image adapted from Hugging Face blog [40]

#### 2.4.3 API call builder

The intent label enables construction of a GDC API call with the required entity parameters using rule-based logic, such as those shown for *ssm frequency* and *freq cnv loss or gain* (Figure 4, 5). In the SSM case, for each SSM, the */ssm occurrences* endpoint is queried using the *ssm id* obtained from the /ssm API endpoint. The */cnv occurrences* endpoint is queried for CNVs. Depending on the intent, different API calls will be constructed and executed.

#### 2.4.4 Query GDC and process results

For frequency of mutation co-occurence, or the fraction of cases that contain a combination of two different SSMs, CNVs, or combinations of SSMs and CNVs, the API call is executed separately for each variant and a joint frequency is computed from the overlapping set of cases and reported as the GDC Result. For single SSMs or CNVs, only a single variant frequency is returned in the GDC Result and involves fewer API calls. (Figure 4, 5).

#### 2.4.5 Generate prompts for the LLM

The GDC Result is appended to the input natural language query to construct a query augmented prompt. A descriptive prompt is also constructed using the gene entities and intent from the query, to obtain a natural language response from the LLM based on genes and intent.

#### 2.4.6 Generate LLM response R

The Guidance library from Microsoft is used to constrain generation with a regex for variant frequency in percentage and structure the LLM response R from llama-3B [33]. Constrained decoding enables easy extraction of variant frequencies from LLM response R (Figures 4, 5). A descriptive response is also obtained using a similar set up, where the LLM will generate a description based on genes and intent in cancer.

#### 2.4.7 Final check and confirmation

In the final step, the percentage in the final LLM response R is checked against the GDC Result to ensure accuracy of frequency, and changed to the GDC Result if inaccurate. This step ensures no additional hallucinations are introduced in the final response. The descriptive response is postprocessed for sentence completion and a natural language response is returned by concatenating the descriptive response with the variant frequency.

In addition to direct testing of the GDC-QAG app via Gradio, the exposed MCP server URL through Gradio can also be integrated with frameworks like LangGraph to programmatically execute the GDC-QAG MCP tool on an input query [34]. The MCP server provides the GDC-QAG MCP tool that is obtained by the MCP client and used to process an input query.

### 2.5 Novel features of GDC-QAG

#### 2.5.1 Accept natural language queries with broad cancer types

In addition to the specific cancer types directly mentioned in the query (see Table 1), GDC-QAG also attempts to decompose broad cancer types mentioned in the query into specific cancer projects in the GDC and report a frequency per-cancer type. For example: “melanoma” in the query “What is the frequency of BRAF V600E in melanoma in genomic data commons?” will be decomposed into twelve different GDC projects ‘TCGA-SKCM’, ‘EXCEPTIONAL RESPONDERS-ER’, ‘MATCH-P’, ‘MATCH-C1’, ‘MATCH-Z1I’, ‘MATCH-R’, ‘HCMI-CMDC’, ‘CPTAC-3’, ‘FM-AD’, ‘TCGA-UVM’, ‘MATCH-S2’, and ‘MATCH-I’ and frequency reported per-cancer in the result. The query “What is the frequency of JAK2 V617F in lymphoid leukemia?” will be decomposed into five GDC projects ‘TARGET-ALL-P1’, ‘TARGET-ALL-P2’, ‘MP2PRT-ALL’, ‘TARGET-ALL-P3’, ‘HCMI-CMDC’ and a frequency reported for each cancer.

#### 2.5.2 Compute frequency of combinations of SSMs and assess impact on survival

In addition to the use cases presented in Table 1, GDC-QAG can compute i) frequency of cases with two different simple somatic mutations, and ii) frequency of cases with a cnv amplification or deep deletion and a simple somatic mutation. This capability can be used to systematically identify new variant combinations that occur in appreciable frequencies in different cancer types, estimate the frequencies of previously published combinations in the GDC, or analyze survival curves for combinations of mutations. For instance, using single variant frequencies from the GDC, we programmatically identified a few combinations of variants that co-occur in a few different cancer types (Table 3). We analyzed survival curves for the low grade glioma example (Figure S7). Using GDC-QAG, we identified cases carrying both mutations, IDH1 R132H and TP53 R273C and analyzed survival curves for the single mutants, double mutants, and wild-type cases that lack mutations in both IDH1 and TP53. The survival curves show that patients with the IDH1 R132H mutation tend to have a median survival time that is roughly 1.6 times greater than the double mutants (median OS=100 months versus 62 months for IDH1 R132H mutation and double mutant respectively, *P <* 6.6*e −* 3 Log Rank Test), suggesting this mutation is beneficial in low grade glioma. These findings are consistent with a previous publication that analyzed missense mutations in TP53 and IDH1 and studied the impact of single and double mutants on overall survival in low grade glioma [35]. Overall this example analysis shows that GDC-QAG can be used to systematically identify single and combination mutations that occur in appreciable frequencies and impact survival in different cancers.

**Table 3:** Natural language questions for mutation combinations. Mutation combinations identified from the GDC where frequency of each single simple somatic mutation is atleast 10% in a specific cancer type, and combinations created and analyzed with GDC-QAG. Results from all models are shown. *GV T* frequencies are obtained using independent functions that directly query the GDC API, see Table 2. Baseline LLM responses for Qwen-4B and llama-3B models are obtained using vLLM==0.7.2 and transformers==4.49.0 versions. GPT-4o model (gpt-4o-2024-08-06) tested using the OpenAI client (Methods).

| Example questions | GDC-QAG | *GVT* | GPT-4o | Qwen-4B | llama-3B |
| --- | --- | --- | --- | --- | --- |
| What is the co-occurrence frequency of IDH1 R132H and TP53 R273C simple somatic mutations in the low grade glioma project TCGA-LGG in the genomic data commons? | 7.02% | 7.02% | 0.5% | 0.46% | 0.01% |
| What is the co-occurrence frequency of RNF43 G659Vfs*41 and RPL22 K15Rfs*5 simple somatic mutations in the TCGA-UCEC project? | 6.45% | 6.45% | 0.12% | 0.07% | 0.0% |
| What is the co-occurrence frequency of ACVR2A K437Rfs*5 and RPL22 K15Rfs*5 simple somatic mutations in the TCGA-STAD project? | 8.76% | 8.76% | 0.12% | 0.68% | 0.0003% |
| What is the frequency of cases with amplifications in EGFR and simple somatic mutations in ATRX in the TCGA-LGG project? | 0.97% | 0.97% | 3.5% | 10.4% | 0.01% |

#### 2.5.3 Reusable pipeline components for other applications

QAG is implemented in a modular fashion such that the individual pipeline components can be reused for other research applications.

The GDC is focused on somatic variants. Often researchers are also interested in obtaining the germline frequencies of the variant for somatic or germline variant classification, or to assess the pathogenecity of variants using other data sources such as gnomAD [36]. The GDC employs various API endpoints such as the /ssm endpoint which can be used in concert with GDC-QAG to infer gene and mutation entities and variant position, which can then be subsequently fed into a gnomAD graphQL query to obtain germline frequencies.

The /ssm API endpoint in the GDC can also be used to obtain a dbSNP rsID which can be input into the litvar2 /variant API endpoint to obtain all pubmed publications (PMIDs) for a variant [37]. This application can be useful to automatically collate the number publications that exist for a variant, useful for variants of unknown significance.

On the therapy front, the CIViC database contains expert curated therapies for a variant. The gene and mutation output of QAG entity extraction can be directly used to construct a graphQL query for CIViC database that returns all therapies for the somatic variant in question.

## 3 Discussion

GDC-QAG is an algorithm that leverages extensive APIs available in the GDC, focusing on open data and metadata. The GDC is the world’s largest platform for cancer research that provides access to around 9.5 PB of high-quality harmonized clinical, genetic, transcriptomic and other types of omics data from 86 different cancer projects. The GDC was one of the first cancer research systems to produce a very rich set of API endpoints to allow a cancer researcher to easily search and retrieve data, including a variety of pre-computed analysis results and endpoints. Such high-quality data in a data commons like the GDC has great potential to improve inferences by large LLMs, to increase trust in the responses and mitigate hallucinations, especially for factual queries in precision oncology centered around numeric facts or variant frequencies.

In this study, we prepared a *GV T* evaluation test set comprising questions that cover a range of precision oncology use cases including corresponding variant frequencies obtained from the GDC. We observe an overall systematic underestimation of variant frequencies in baseline LLM responses compared with *GV T* , with some use cases more affected than others. Different LLMs appear to perform better at different tasks. For instance, the Qwen-4B model performs better than GPT-4o and llama-3B on gain and amplification queries. It is also interesting that a large model like GPT-4o does not perform better than smaller models like Qwen-4B and llama-3B on our evaluation dataset, suggesting external context needs to be supplied for numeric information such as variant frequencies irrespective of model size, particularly given zero shot prompting. We found variant frequencies from GDC-QAG and *GV T* to be entirely consistent, suggesting greatly improved results can be obtained using GDC-QAG on precision oncology queries for the use cases presented in this study.

Our approach has several limitations. Disease recognition from the query currently relies on a combination of direct matches to GDC project descriptions and initial cancer entity guess using small NLP models. Gene and mutation entity recognition is also restricted to protein mutations and focused on string search using data from the GDC. Further enhancements could be achieved by developing and evaluating named entity recognition models, as well as by continuing the training of small NLP models. The current version of GDC-QAG can also only return frequencies for two variants in combination for one cancer type at a time, but generalization to triples and higher-order combinations is straightforward.

The other limitation is that GDC-QAG is currently use-case driven, with templated use cases implemented focusing on molecular mutational profiles. Any use cases outside of the templates may not work well. Our approach will need to be extended to accomodate more templates and biological use cases across modalities such as estimating frequencies of variants from both DNA and RNA, frequencies of fusions and copy number variants from both DNA and RNA data, and other questions that incorporate multi-modal combinations. To do so, the BERT model will need to be re-trained on paired queries and intent labels, and the parameters of the new API calls programmed into the code for execution. While the current framework is suited for specific research tasks, to generalize across queries and use cases, open-source LLMs will need to be trained on paired ground-truth datasets of natural language queries and API calls in an agentic framework. Trained agents have the potential to simplify querying across modalities, leading to unique biological discoveries in data meshes and other multi-modal ecosystems that expose APIs ([38, 39]).

## 4 Figures and Tables

**Supplementary Figure S1:**
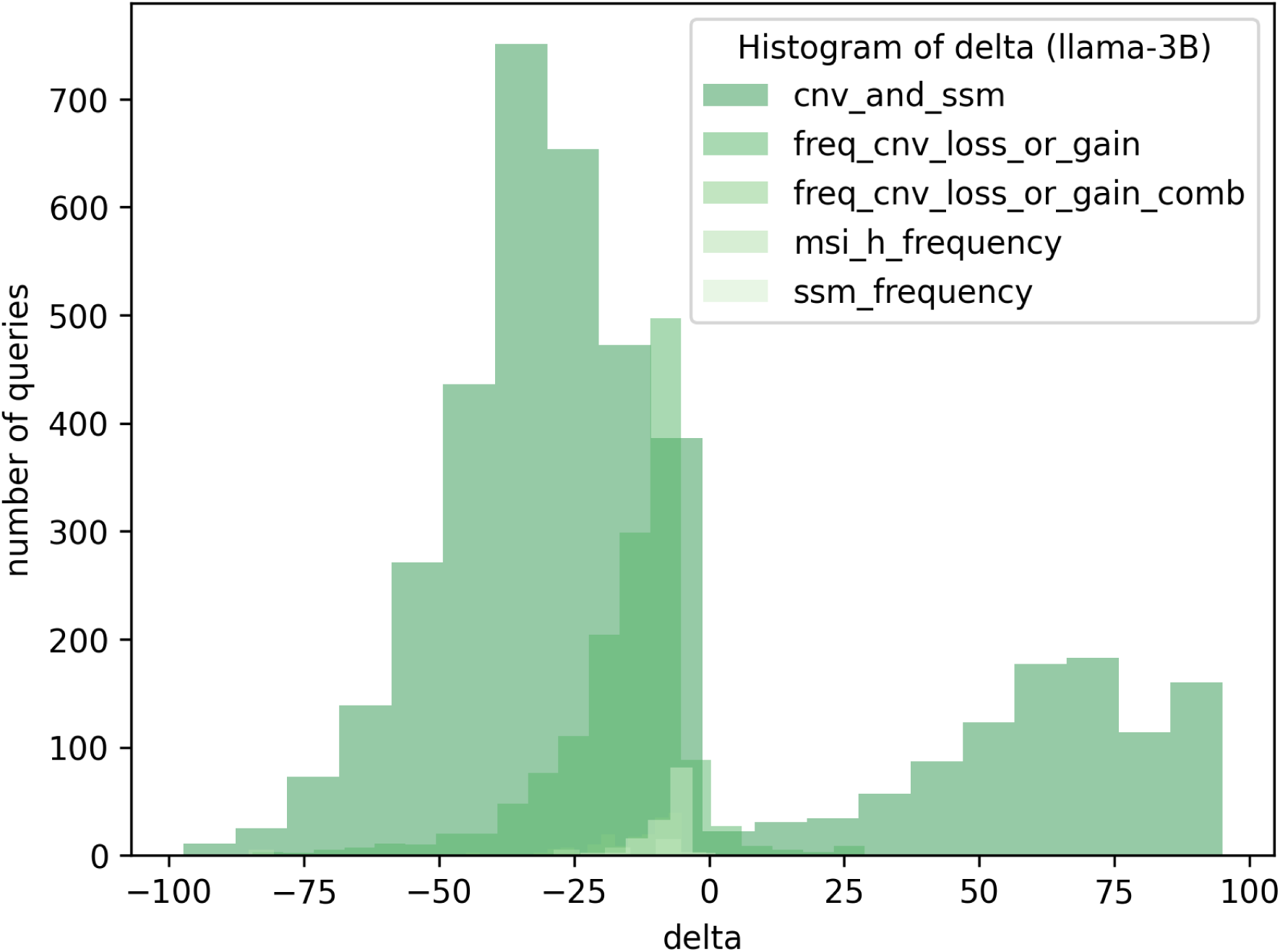
Difference between llama-3B and *GV T*

**Supplementary Figure S2:**
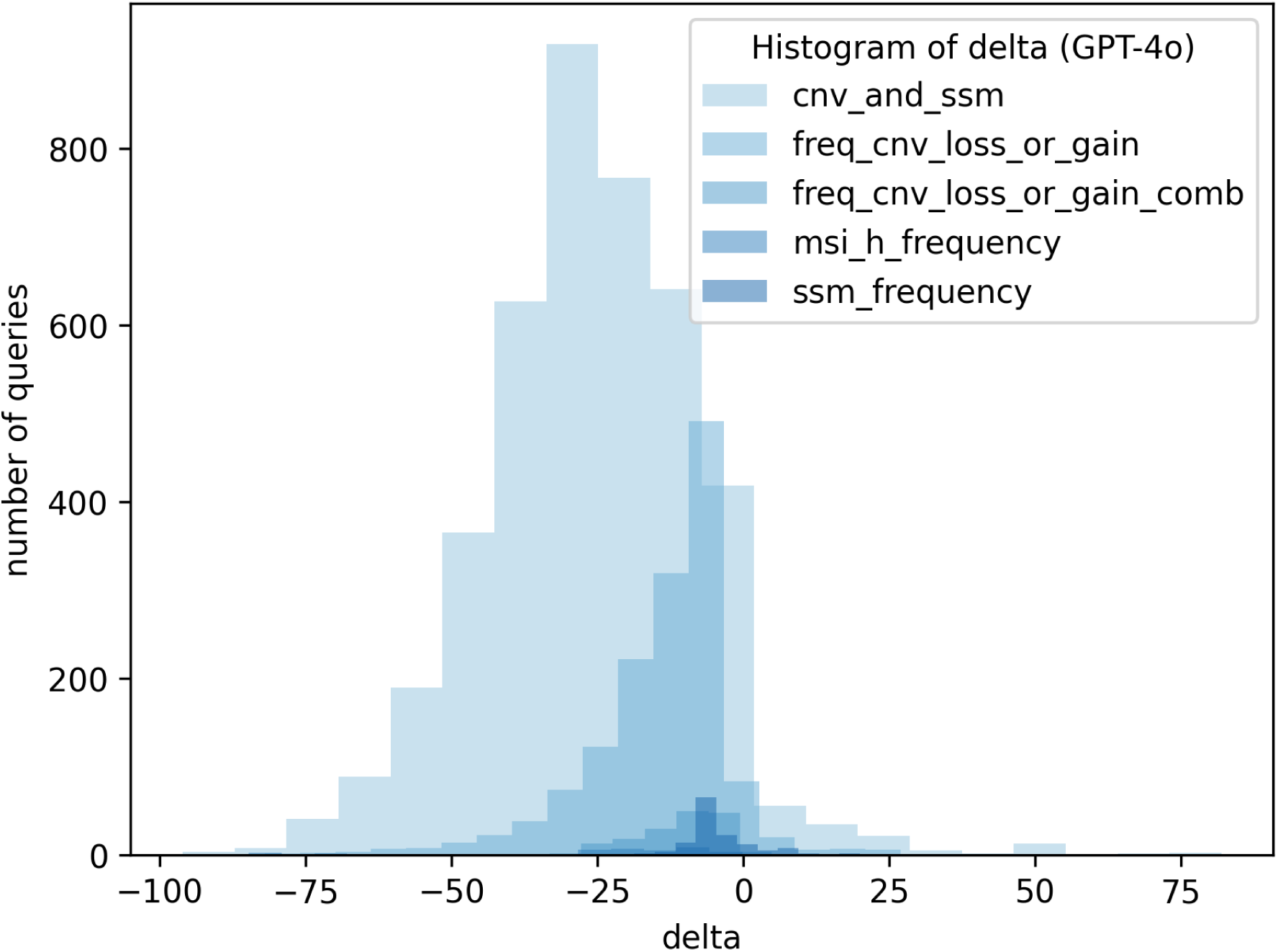
Difference between GPT-4o and *GV T*

**Supplementary Figure S3:**
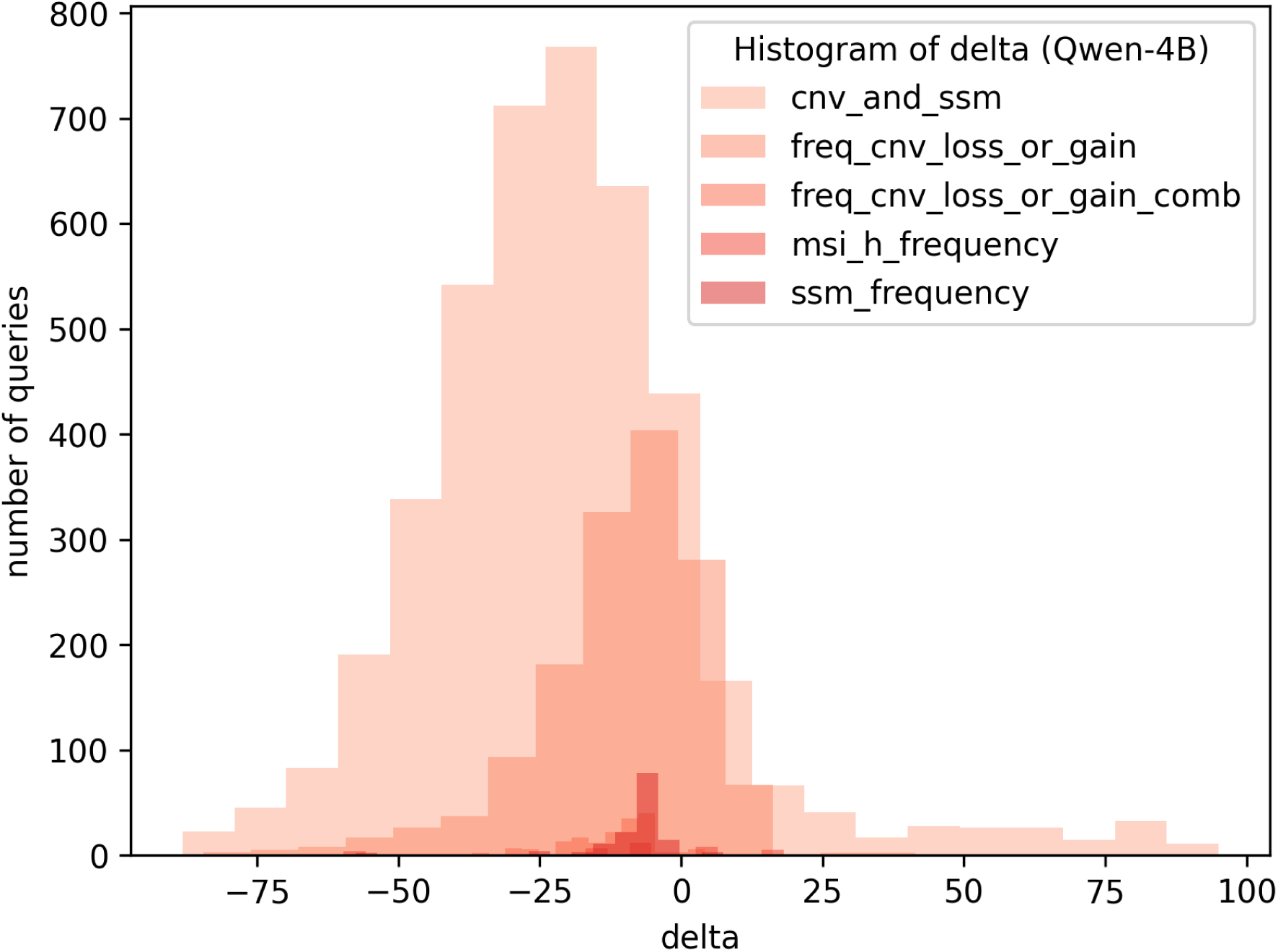
Difference between Qwen-4B and *GV T*

**Supplementary Figure S4:**
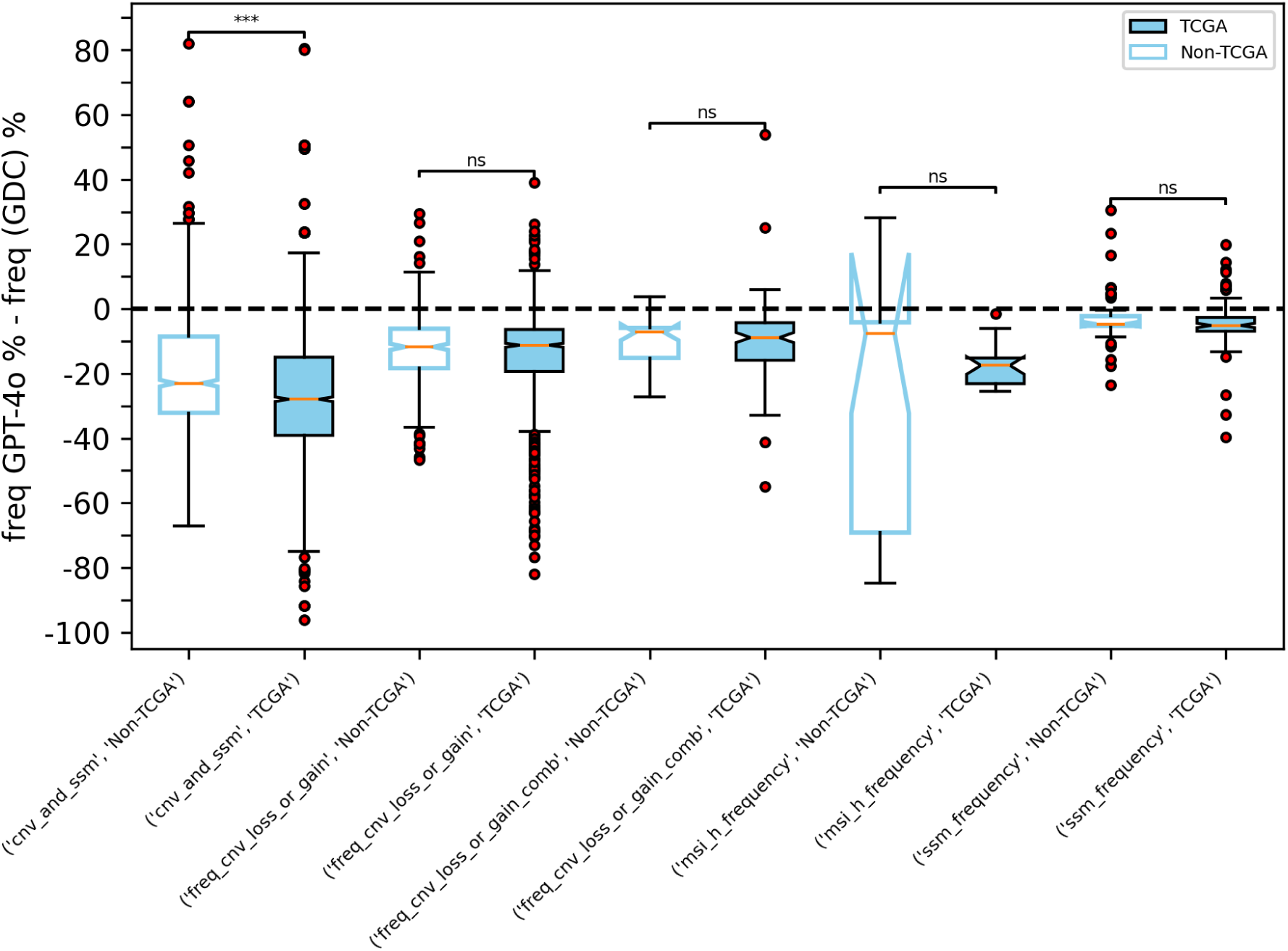
Comparison of GPT-4o model performance by TCGA and non-TCGA questions. Significance calculated using t-tests. **\*\*\*** *P <* 0.001, **\*\*** *P <* 0.01 , **\*** *P <* 0.05, **ns** *non − significant*

**Supplementary Figure S5:**
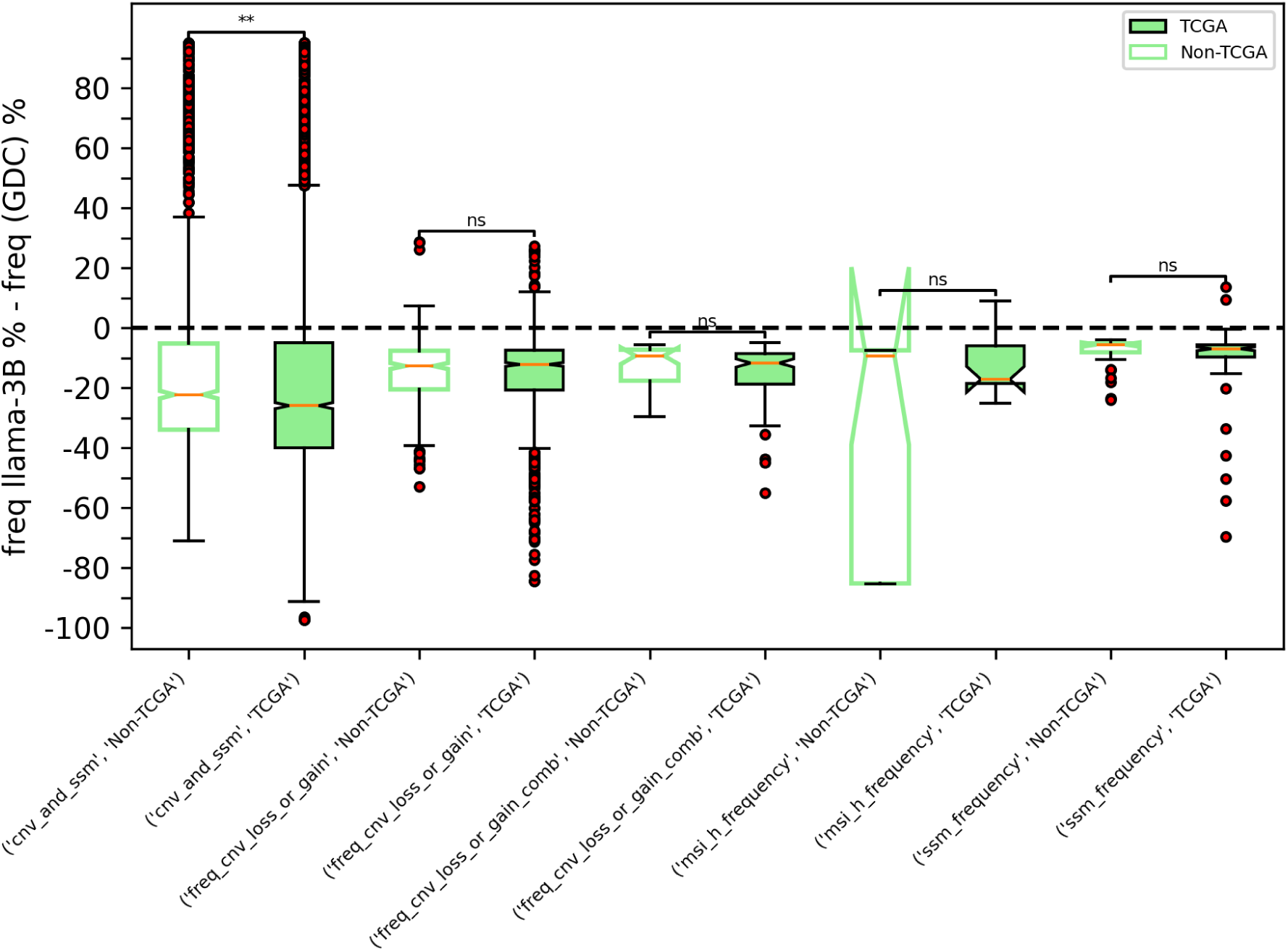
Comparison of llama-3B model performance by TCGA and non-TCGA questions. Significance calculated using t-tests. **\*\*\*** *P <* 0.001, **\*\*** *P <* 0.01 , **\*** *P <* 0.05, **ns** *non − significant*

**Supplementary Figure S6:**
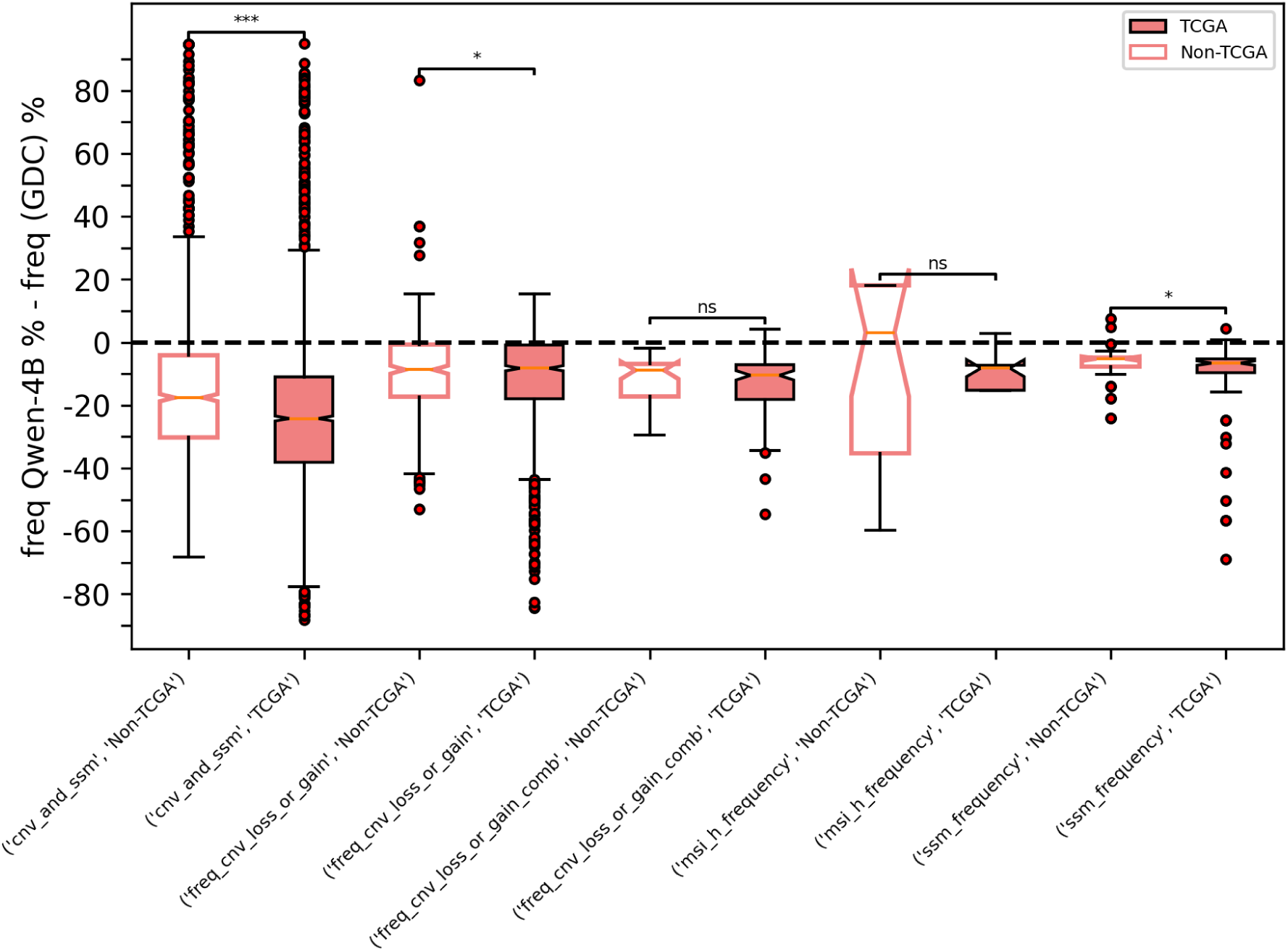
Comparison of Qwen-4B model performance by TCGA and non-TCGA questions. Significance calculated using t-tests. **\*\*\*** *P <* 0.001, **\*\*** *P <* 0.01 , **\*** *P <* 0.05, **ns** *non − significant*

**Supplementary Figure S7:**
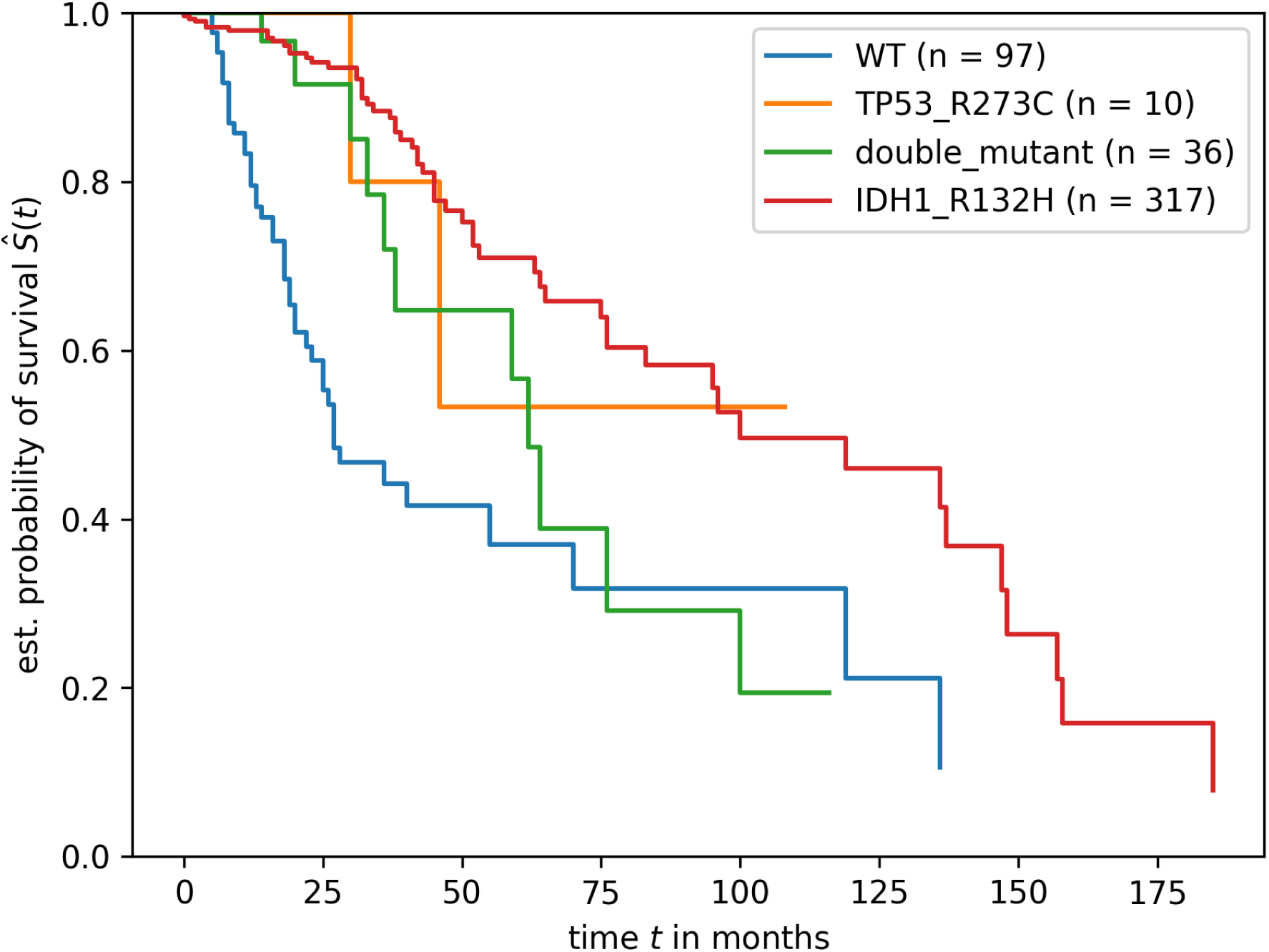
Survival curves for single and double mutants and wild-type (WT) in low grade glioma. The double mutant set contains cases with both TP53 R273C and IDH1 R132H. The single mutant set (TP53 R273C, IDH1 E132H) contains only cases with single mutations (subtracting double mutants). The wild type set is all cases except those with mutations in IDH1 and TP53. Log rank-test is significant for the double mutant and IDH1 R132H comparison. **\*\*\*** *P <* 6*e −* 3

## 5 Methods

### 5.1 Table S1, ChatGPT manual investigation

We constructed a modified prompt where each of the six queries in Table S1 are appended with instructions and tested in the ChatGPT UI powered by GPT-4o, released by OpenAI in May 2024 (gpt-4o-2024-08-06). For an example query, the following is the complete prompt:

What is the frequency of KRAS G12D mutation in the colon adenocarcinoma project (TCGACOAD) in the genomic data commons? Only use results from the genomic data commons in your response and provide frequencies as a percentage. Only report the final response.

### 5.2 Obtaining baseline LLM responses on ***GV T***

For llama-3B and Qwen-4B baseline evaluations, the vLLM framework was used along with sampling with constrained decoding in the llm.generate function to generate the final response, with the following parameters chosen for reproducibility: n=1, temperature=0, seed=1042,max tokens=1000, repetition penalty=1.2 [41]. Guided decoding was set to:

`guided_decoding_params = GuidedDecodingParams (regex=“The final answer is: \d*\.\d*%”)`

We asked the models to only report frequencies from the GDC by appending each query with the following prompt:

Only use results from the genomic data commons in your response and provide frequencies as a percentage. Only report the final response.

A small fraction of test queries 0.5% (n=34/6011) had no reported percentages in the Qwen-4B model responses.

Computations were performed using vllm==0.7.2 and transformers==4.49.0 versions for llama-3B and Qwen-4B baseline evaluations. Fixing the versions of vllm and transformer libraries, together with the parameters listed above maximizes reproducibility, although some limited variability can still be observed in the responses.

GPT-4o was evaluated using the batch API from OpenAI with the following parameters for reproducibility: temperature=0, seed=2000. The prompt was also modified slightly to also return references in the response. Each query was appended with the following prompt to construct the full prompt:

Only use results from the genomic data commons in your response and provide frequencies as a percentage in the result. Report the result in the following output JSON format, strictly using the structure:

{

`“result”: “The final answer is: frequency %”,`

`“references”: “list of references”`

}

For GDC-QAG evaluation, each query is augmented with the following prompt and passed to the llama-3B model along with frequency information retrieved from the GDC:

Only report the final response. Ignore all prior knowledge. You must only respond with the following percentage frequencies in your response, no other response is allowed: < GDCResult >

Occasionally the GDC API connection aborts due to remote disconnection, in which case the query is re-tried to resolve the issue.

Results for Table 3 are also obtained using the same approaches.

### 5.3 Curation of ***GV T*** for baseline LLM evaluations

#### 5.3.1 CIViC dataset and template question design

We used the clinical evidence nightly summaries, a TSV file that is downloadable from the CIViC website, that contains clinical evidence statements for each variant with trustable medical literature. Considering variants where the gene name and variant change in the protein are fully specified (e.g. BRAF V600E as opposed to BRAF V600), and excluding fusions and frameshift mutations, we constructed a filtered CIViC molecular profile (n=699) (Table S2). We then defined 24 template synthetic questions focused on simple somatic mutations, copy number variants and MSI, with placeholders added for genes, mutations and cancer type (Table S3). These template questions were manually curated, and some natural language diversity was introduced using GPT-4o (gpt-4o-2024-08-06).

#### 5.3.2 Queries focused on combination of copy number variants

To design queries for combinations of copy number variants, we used the results from Figure 3 of the Cheng et al publication focused on homozygous and heterozygous deletions spanning 9 unique gene combinations [31]. The gene combinations included ‘CDKN2A’ and ‘PTEN’, ‘NF1’ and ‘BRCA2’, ‘RB1’ and ‘CYLD’, ‘FAT1’ and ‘MAP2K4’, ‘TET1’ and ‘CDH1’, ‘TP53’and ‘SMARCB1’, ‘BIRC2’ and ‘CDKN2C’, ‘BIRC3’ and ‘CDKN2C’. The limited set of 18 pancancer queries were designed using a standard template (Table S4).

#### 5.3.3 Curation of synthetic questions using CIViC, GDC and publication data

For each row in the filtered CIViC molecular profile, we sampled n=5 template questions at random, and substituted the gene, mutation and disease placeholders in the template question with the corresponding data from the filtered CIViC molecular profile (699*5=3495). Dropping duplicate questions, each of the n=1822 unique questions had a broad cancer type substituted under the ”disease” place-holder (e.g melanoma), as opposed to a specific cancer project in the GDC (e.g. TCGA-SKCM) since disease information is obtained from the CIViC database (see “disease” column in Table S2).

To design GDC project-specific questions where variants have a frequency of at least 5% in the GDC, we performed an initial run of the 1822 questions through GDC-QAG which decomposed the broad cancer type in the query into specific cancer projects in the GDC. For instance, for the following melanoma query:

What is the frequency of MSI-H in Melanoma in the genomic data commons?

12 cancer projects were returned by GDC-QAG: ‘TCGA-SKCM’, ‘EXCEPTIONAL RESPONDERSER’, ‘MATCH-P’, ‘MATCH-C1’, ‘MATCH-Z1I’, ‘MATCH-R’, ‘HCMI-CMDC’, ‘CPTAC-3’, ‘FM-AD’, ‘TCGA-UVM’, ‘MATCH-S2’, and ‘MATCH-I’, and a frequency obtained for each project. If a disease could not be mapped to a cancer project by GDC-QAG, all 86 GDC projects were returned and queried. The initial set of 1822 queries were then reformatted using a custom script to include GDC project-specific information. Only those queries that returned a ground-truth variant frequency of at least 5% were retained in *GV T* .

Questions around combination copy number variants (Table S4) were also run through GDC-QAG and queries reformatted to contain GDC project-specific information for disease. Similar to above, only those queries that returned a ground-truth variant frequency of at least 5% were retained in *GV T* .

To increase coverage of simple somatic mutations in *GV T* , we also downloaded a TSV of all simple somatic mutation frequencies from the GDC using the mutation tab in the mutation frequency app, and constructed additional template synthetic questions using the same approach as described for CIViC data.

#### 5.3.4 Generation of ***GV T*** variant frequencies

Variant frequencies (truth) were generated for *GV T* by dividing the n=6011 synthetic questions into seven different template categories (cnv and ssm, heterozygous and homozygous deletions, gains, amplifications, ssm frequency, msi frequency, and frequency of CNV losses or gains (combinations)) and writing independent functions that query the GDC API using entities extracted from the query. See Table 2 for API endpoints and parameters used for the different query types. The final *GV T* dataset with n=6011 questions and variant frequencies is available in Table S5.

### 5.4 Entity extraction: Gene and Protein mutations

To preconstruct the GDC gene-mutation dictionary we queried the /ssms API endpoint using *fields* = *gene aa change* and downloaded the ssm data from the API in batches. We then loaded the data in a dictionary with genes as the keys and protein mutations as the values. The pre-constructed dictionary contains n=19606 genes and 5.6 million ssms scraped from the GDC /ssms API endpoint.

The dictionary is available as a dataset in Hugging Face

https://HuggingFace.co/datasets/uc-ctds/GDC-QAG-genes-mutations

### 5.5 Entity extraction: Cancer entities

Using the results of the /projects API endpoint, we first search for a GDC project-id in the query that maps to a unique project, and consider those entities if a match is found. If not, initial guesses for cancer entities are made using two scispacy NLP models from allenai [32]: i) bc5cdr model trained on the bc5cdr corpus, a biocreative disease relations dataset that is an annotated dataset of chemicals, diseases and interactions in 1500 pubmed articles, and ii) the encore websm model for medical data with larger vocabulary. These models recognize broad entities such as lymphoid leukemia, myeloma, or breast cancer from the query. We then postprocess these initial guesses in different ways to infer GDC project-ids: i) using a wildcard match for the initial cancer entity in the /projects endpoint to return a project-id, ii) matches between query terms and project name and descriptions from the /projects API endpoint. If no cancer entities are obtained using these described approaches, all projects from the GDC are returned in GDC-QAG and used for querying variant frequencies per project. GDC-QAG relies on GDC /project endpoint to accurately map disease information from a query to a specific GDC project. For example, “breast invasive carcinoma” mentioned in the query will map the query to the ‘TCGA-BRCA’ project.

### 5.6 Intent inference

The GDC-QAG algorithm performs user intent inference to map the query to an appropriate GDC API endpoint. This step involves predicting a user intent label from a query. The following user intent labels are supported:

`ssm_frequency`

`msi_h_frequency`

`freq_cnv_loss_or_gain`

`cnv_and_ssm`

`top_cases_counts_by_gene`

The *top cases counts by gene* intent label also maps broadly to the *cnv and ssm* use case, with the GDC API endpoint returning the number of cases with CNVs or SSMs or both. We defined starting template questions for each user intent. Example questions for intents *ssm frequency*, *msi h frequency* and *freq cnv loss or gain*, respectively, are shown below:

What percentage of [cancer] cases have the [gene] [mutation] mutation?

What is the frequency of microsatellite instability-high in [cancer]?

What is the frequency of somatic [gene] heterozygous deletion in [cancer]?

To generate natural language diversity for BERT classifer training, we asked PhonenixAI, a wrapper tool around GPT-4o (gpt-4o-2024-08-06) to generate multiple variations of each template question, and collated n=126 template questions in total for five user intents. We then queried the */ssms* GDC API endpoint and obtained the top 1000 results with genes, mutations and cancer information and filtered this set to include fully specific SSMs (excluded fusions and other unspecified variant changes) (n=506). We then generated n=63756 labeled questions (506*126) and intent pairs by looping through the set of template questions and filling the placeholders using information from the GDC. This dataset was used to train a *bert-base-uncased* model, with a train:validation split of 70:30 created using the sklearn package and trained for two epochs using PyTorch. Model performance was tested on an independent test dataset with 20428 questions assembled from the /ssms endpoint with completely different genes and mutations observed in different cancer types. Accuracy, precision, recall and F1 score for all tested intents were 1.0, assessed using the classification report and confusion matrix packages from sklearn. The entire test dataset used for testing model performance, including the prediction output, is available in Table S10.

The entire dataset used for training is available in Hugging Face

https://HuggingFace.co/datasets/uc-ctds/query_intent_dataset

The trained BERT classifier model (intent model) is also available in Hugging Face, with the following model id:

uc-ctds/query_intent

### 5.7 GDC API endpoints used for various computations

#### 5.7.1 Frequency of simple somatic mutations or SSMs

For SSM frequency, the /ssms endpoint is used to query by gene and protein mutation and obtain an ssm id which is then input into */ssm occurrences* endpoint to get the list of cases with the SSM for the cancer project in question. Counts are then converted into frequencies by obtaining the number of cases with available SSM data in the cancer project from the */case ssms* endpoint. For combination SSMs, the result for each SSM is obtained separately and the intersect of cases is calculated to obtain the joint frequency of cases carrying both mutations for the cancer project mentioned in the query. The result from the computations are formatted into a one-line helper result (GDC Result) and passed to the prompt for the llama-3B model in GDC-QAG.

#### 5.7.2 Frequency of CNV loss or gain

For frequency of CNV loss or gain a set of loss terms: ‘loss’, ‘LOH’, ‘deletion’, ‘co-deletion’, ‘homozygous’ are defined to match with the query and obtain the right cnv change type and change 5 category for constructing the GDC API parameters. Gains and amplifications are defined using the same logic. For each cancer and gene entity obtained from the query, the */cnv occurrences* endpoint is queried using the parameters cnv change, cnv change 5 category, gene symbol and GDC project id to obtain the list of cases with the CNV. Counts are converted into frequency by obtaining the number of cases with available CNV data using the */case ssms* endpoint. For combination CNVs spanning multiple genes, the result for each gene and CNV change type is calculated separately and the intersect of cases is computed and converted to the joint frequency using the */case ssms* endpoint. The result from the computations are formatted into a one-line helper result (GDC Result) and passed to the prompt for the llama-3B model in GDC-QAG.

#### 5.7.3 Frequency of MSI

For MSI frequency, the frequency of BAM files with MSI tag is computed for the required cancer project and considered as the MSI frequency. The /files endpoint is queried using the GDC project id and the following fields: data format BAM, experimental strategy WGS,WXS. We only score tumors where MSI status is reported from the BAM files, and exclude any null inferences. MSI score can be either ‘MSS’ or ‘MSI’ for microsatellite stable or unstable, respectively. Frequency of MSI is calculated by dividing the number of BAMs with ‘MSI’ assigned for MSI score by the total number of cases (‘MSI’ + ‘MSS’). The result from the computations are formatted into a one-line helper result (GDC Result) and passed to the prompt for the llama-3B model in GDC-QAG.

#### 5.7.4 Frequency of CNV and SSMs

To calculate the number of cases with CNVs and/or SSMs, the */top cases counts by genes* endpoint is queried using the gene entity (Ensembl gene id) mentioned in the query. This endpoint returns the number of cases with mutations (CNVs or SSMs, or both). Total number of cases with variation data for the cancer project is obtained using the */case ssms* endpoint where available variation data is set to both CNVs and SSMs. Counts are then converted to frequency and reported in the result. For queries that mention both deletion and an SSM, or amplification and an SSM, the frequency of CNV loss or gain and frequency of SSMs are calculated separately as described in the previous sections, and an overlap of cases with both SSM and CNV is calculated by computing the intersect. In the current implementation, the natural language queries should be formulated to mention the CNV type first (e.g. amplification or deletion) followed by SSMs. A joint frequency is then obtained by dividing the number of cases in the intersect with the total variation data available for the project. The result from the computations are formatted into a one-line helper result (GDC Result) and passed to the prompt for the llama-3B model in GDC-QAG.

### 5.8 Survival curves, Fig. S4

Using GDC-QAG the case ids of double mutants with both IDH1 R132H and TP53 R273C mutations in TCGA-LGG were retrieved. A double mutant cohort was created by uploading the *case ids* in the GDC portal. Single mutant cohorts were created using the GDC portal, subtracting cases with double mutants. The wild-type cohort was also created using the GDC portal, excluding cases with mutations in IDH1 and TP53. Clinical data for LGG was downloaded from the portal. Survival curves were generated for pairwise combinations to confirm trends with previous publication (WT, TP53 R273C) and (double mutant, IDH1 R132H), and survival plot data downloaded and combined to generate Fig. S4.

### 5.9 Gradio deployment in Hugging Face Spaces

The baseline LLM evaluations including constrained decoding were obtained using vllm==0.7.2 and transformers==4.49.0 versions, tested using the gdc-qag GitHub code on a single V100 GPU node with 16GB GPU RAM on our on-prem GPU cluster. To deploy on Hugging Face and run on a slice of H200 ZeroGPU, the vllm functions were changed to the Transformers equivalent, and constrained decoding was implemented using the Guidance library from Microsoft. The GDC-QAG functionality is wrapped using Gradio and exposed as an MCP server on Hugging Face spaces. Note that the guidance generation can differ from vLLM generation.

The app is available for testing through Hugging Face ZeroGPU, accessible at

https://HuggingFace.co/spaces/uc-ctds/GDC-QAG

## 6 Data availability

This project uses open-access data available in the GDC that is queried using the various API endpoints. The results of the paper are based on GDC data release (DR) 43, May 7 2025 version of the GDC.

## 7 Software availability

GDC-QAG is freely available via GitHub, including the code to reproduce the figures

https://github.com/uc-cdis/gdc-qag

The GDC-QAG app can be tested using Hugging Face ZeroGPU, accessible at:

https://huggingface.co/spaces/uc-ctds/GDC-QAG

## 8 Supplementary Data

Supplementary figures and tables are available in the excel file.

## Acknowledgements

This work was funded in part through the Advanced Research Projects Agency for Health (ARPA-H) under contract 75N92020D00021/5N92023F00002. The views and conclusions contained in this document are those of the authors and should not be interpreted as representing the official policies, either expressed or implied, of the U.S. Government.

## 9 Author information

### 9.1 Authors and Affiliations

#### 9.1.1 Center for Translational Data Science, University of Chicago, Chicago IL

Aarti Venkat, Steven Song, Anirudh Subramanyam, Michael Lukowski, William P. Wysocki, Zhenyu Zhang, Robert L. Grossman

#### 9.1.2 Section of Biomedical Data Science, Department of Medicine, University of Chicago, Chicago IL

Aarti Venkat, Robert L. Grossman

#### 9.1.3 Department of Computer Science, University of Chicago, Chicago, IL

Steven Song, Michael Lukowski, Robert L. Grossman

#### 9.1.4 Medical Scientist Training Program, Pritzker School of Medicine, University of Chicago, Chicago, IL

Steven Song

### 9.2 Contributions

A.V, S.S, R.L.G. contributed to the study design, experiments and interpretation. A.V. drafted the manuscript with critical revisions and comments provided by S.S. and R.L.G. and the other authors. A.S. contributed to the methods and interpretation, including app development on Hugging Face. M.L. contributed to app deployment. Z.Z. and W.P.W. reviewed GDC API calls and integration. W.P.W. wrote code for independent validation of GDC-QAG frequencies and *GV T* variant frequencies. All authors read and contributed to manuscript revisions and approval of the final version.

### 9.3 Corresponding author

Correspondence to Aarti Venkat

